# Sustained ON alpha retinal ganglion cells in the temporal retina exhibit task-specific regional adaptions in dendritic signal integration

**DOI:** 10.1101/2024.03.27.586958

**Authors:** Jonathan Oesterle, Yanli Ran, Paul Stahr, Jason ND Kerr, Timm Schubert, Philipp Berens, Thomas Euler

**Author notes:** These authors contributed equally (Jonathan Oesterle, Yanli Ran).

## Abstract

Various retinal ganglion cells (RGCs) show regional adaptations, for instance, to increase visual acuity. However, for many RGC types, it is not known how they vary in their signalprocessing properties across the retina. In the mouse retina, sustained ON alpha (sONα) RGCs were found to have differences in morphology and receptive field sizes along the nasotemporal axis, and temporal sONα RGCs are likely to play a role in visually guided hunting. Thus, we hypothesised that this cell type also exhibits regional adaptations on the level of dendritic signal processing and that these adaptations are advantageous for prey capture. Here, we measured dendritic signals from individual sONα RGCs at different locations in the *exvivo* whole-mount mouse retina using two-photon microscopy. We measured both postsynaptic Ca^2+^ signals at the dendrites of individual RGCs and presynaptic glutamate signals from bipolar cells (BCs). We found that temporal sONα RGC dendrites exhibit, in addition to the expected sustained-ON signals with only weak surrounds, signals with strong surround suppression, which were not present in nasal sONα RGCs. This difference was also present in the excitatory presynaptic inputs from BCs, suggesting a presynaptic origin. Finally, using population models in an encoder-decoder paradigm, we showed that these adaptations might be beneficial for detecting crickets in hunting behaviour.

## Introduction

The architecture of the visual system of animals is shaped by the statistics of the environment as well as behavioural demands (1). Thus, although their retina is based on a common blueprint, vertebrates show significant variations in retinal architecture, including many regional adaptions within the retina. This underscores the influence of evolutionary pressures and ecological niches on visual systems (1, 2).

Some species, like many primates and certain birds, have developed foveas, that is, regional specialisations for high-acuity vision with distinct architecture compared to the peripheral retina (3). But also non-foveated species typically feature local specialisations of their retinas: For instance, zebrafish have a region of higher retinal ganglion cell (RGC) density, also referred to as the “strike zone”, which contains many UV-sensitive photoreceptors and is believed to play a crucial role in hunting (4–6). Similarly, in mice, regional adaptations can already be found at the photoreceptor layer. For example, in some species of the genus *Mus*, including steppe mice (Mus spicilegus) and also the derivative lab strain C57BL/6J, short-(S-) and medium (M-) wavelength-sensitive opsin expression follows a pronounced gradient along the dorso-ventral axis (7–10), resulting in a green-sensitive dorsal retina and a UV-sensitive “hotspot” in the naso-ventral retina (11). These spectral sensitivity differences are propagated via the bipolar cells (BCs) (12, 13) to the RGCs (14). Other mouse species from distinct habitats, such as the wood mice (Apodemus sylvaticus) that—in contrast to steppe mice—are typically found in forests (15), completely lack this gradient (7). At the level of RGCs, mice have also been shown to exhibit a region of lower RGC density in the dorsal retina (16, 17) and several regional adaptations that are specific to distinct RGC types (18, 19).

Here, we focused on sONα RGCs (EyeWire: 8w; (20, 21)), which have been shown to vary across space at the level of their morphology: Temporal sONα have much smaller dendritic arbours and exhibit a higher cell density compared to nasal cells (17, 22, 23). Notably, temporal sONα RGCs have also been linked to visually guided hunting, suggestive of a direct connection between their morphology and functional significance (24, 25).

To better understand if these cells also display adaptations on the functional level and how these arise from their dendritic input and cellular computations, we recorded dendritic calcium signals and excitatory synaptic inputs to sONα cells in different regions of the retina. Based on morphological and functional data, we then created computational population models of both nasal and temporal sONα RGCs. We used these models to encode the visual scene as seen by freely moving mice hunting crickets (25). We trained a decoder to estimate the presence of a cricket from the population responses in a binary classification task. We found that the decoder performed much better for temporal sONα RGCs compared to nasal ones. Moreover, our simulation indicated that stronger surround inhibition already at the level of presynaptic neurons was likely the cellular mechanism responsible for the better performance of temporal sONα RGCs in this task.

Taken together, our results suggest that regional changes in presynaptic circuits and dendritic signal integration are key mechanisms in tuning temporal sOα RGCs for detecting small objects such as insects.

## Results

### Recording sONα RGCs across the retina

To analyse regional adaptations in dendritic signal processing of sONα RGCs, we recorded dendritic Ca^2+^ signals in response to visual stimulation of individual RGCs in the *ex-vivo*, wholemount mouse retina using two-photon imaging. For this, we injected individual RGCs with the fluorescent Ca^2+^ indicator dye Oregon Green BAPTA-1 (OGB-1) using sharp electrodes (Methods), resulting in labelling of individual cells (Fig. 1A, B and Fig. S1). After the functional recordings, we 3D-reconstructed the respective RGCs (Fig. 1) and mapped the dendritic recording field onto the morphology (Methods). We grouped the recorded sONα RGC into three groups based on their retinal location: nasal (*n*) dorsal (*d*), and temporal (*t*) (Fig. 1C). Note that we included the ventral cell in the *n* group because its morphology and functional properties matched this group. We estimated receptive fields (RFs) from a binary dense noise stimulus (20 × 15 pixels, 30 *μ*m per pixel) that was centred on the respective recording fields. As we focused on dendritic recordings, we did not record from RGC somata, but for most cells, from dendrites very close to the soma. These RFs can be used as a proxy for somatic RFs (Fig. S2). Consistent with previous reports (22), we found that, compared to nasal or dorsal cells, temporal sONα RGCs both smaller *dendritic* fields (*n* vs. *d*: p=0.34; *d* vs. *t*: p=0.036; *n* vs. *t*: p=0.0022; Fig. 1D), and smaller (soma-like) *receptive* fields (*n* vs. *d*: p=0.34; *d* vs. *t*: p=0.16; *n* vs. *t*: p=0.013; Fig. 1E), but larger relative RF sizes, i.e. the RF size divided by the dendritic field size (*n* vs. *d*: p=0.89; *d* vs. *t*: p=0.018; *n* vs. *t*: p=0.0089; Fig. 1F). Note that in Fig. 1E, the difference between dorsal and temporal was not significant, likely because of the smaller sample size and one outlier RF in the*d* group (Fig. S2E).

**Fig. 1.**
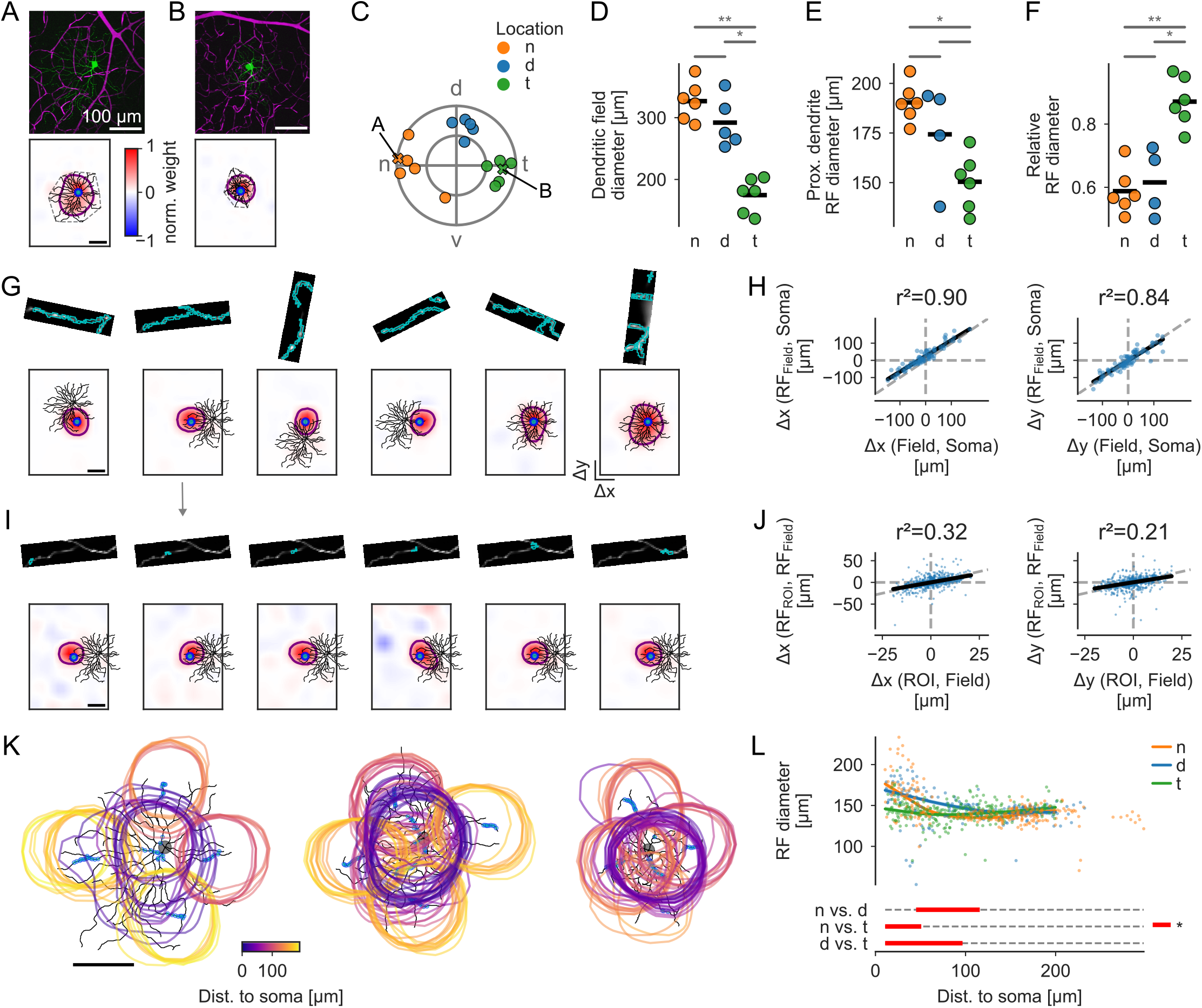
Dendritic receptive fields (RFs) of sONα cells differ between retinal l ocations. **A**. *Top:* Two-photon image of dye-injected nasal sONα RGC (green, OGB-1; magenta, Sulforhodamine 101 (SR-101); weighted two-channel z-projection). *Bottom:* Dendritic skeleton reconstructed from z-stack, its convex hull (grey dashed) overlaid with the RF estimated from a Ca^2+^signal in a proximal dendrite ROI (blue; see Methods), and the respective RF outline estimate (purple ellipse). **B**. As in (A), but for a temporal cell. **C**. Retinal cell locations of all sONα RGCs from which Ca^2+^signals were recorded (*n*, nasal, orange; *d*, dorsal, blue; *t*, temporal, green). RGCs in (A) and (B) are highlighted. **D-F**. Statistical comparison of cells in C, using Kruskal-Wallis and Dunn’s tests with Benjamini-Hochberg correction. **D**. Dendritic field d iameter e stimated f rom c onvex h ull (see M ethods). **E**. P roximal d endrite R F d iameter, i f it was recorded. **F**. Proximal dendrite RF diameter divided by dendritic field d iameter. **G**. Morphology of RGC in (A), overlaid with ROIs and respective RFs. *Top row:* Ca^2+^signal of dendritic recording fields averaged over time with highlighted ROIs (blue). *Bottom row:*Morphology (black) and location of respective recording fields (blue) o verlaid w ith o utline (purple) o f fi eld RF (M ethods). **H**. Linear regression and *r*^2^ from Pearson’s Correlation for field RF centre positions w.r.t. soma versus field centre positions w.r.t. to soma for x (*left*) and y (*right*), respectively. **I**. *Top row:* As in (G), but for RFs from individual ROIs from the second field in G. *Bottom row:* As in (G), but for the respective ROI RF. **J**. Robust linear regression and *r*^2^ from Percentage Bend Correlation for ROI RF centre w.r.t. Field RF centre versus ROI centre w.r.t. field centre for x (*left*) and y (*right*), r espectively. **K**. All ROIs and ROI RF outlines after quality filtering (Methods; RF outlines colour code ROI distance to soma), for a *n*, a *d* cell, and a *t* cell, respectively. **L**. Diameters of ROI RF outline estimate (Methods) as a function of ROI distance to soma by retinal region (colour), and fits from a Generalised Additive Model (see Methods). Distances of significant difference between the groups are highlighted (bottom; red).

### Dendritic signals reflect localised input processing

Next, we measured Ca^2+^ signals across the dendritic arbour of individual cells. For each dendritic recording field (32 × 16 pixels, 31.25 Hz), we extracted regions-of-interest (ROIs) using local pixel correlations (Methods). We estimated an RF using the aforementioned dense noise stimulus for each ROI and additionally for each field by combining the respective ROIs in each field (Methods). The RF centres followed the location of the dendritic recording fields across the dendritic trees (Fig. 1G, H). Even within fields, the relative RF centre positions (w.r.t. the field RF) were correlated with the relative position of the individual ROIs (w.r.t. the field centre) (Fig. 1I-K), suggesting that the recorded signals were electrically isolated dendritic signals and not mainly back-propagated somatic signals.

In dendrites close to the soma, *n* cells and *d* cells had larger RFs than *t* cells, with no significant difference between *n* and *d* cells (Fig. 1K, L). However, for more distal dendrites (≥ 116 *μ*m) there were no significant differences between the retinal locations (Fig. 1K, L). This suggests that the number of excitatory inputs on different retinal locations is likely comparable between *n* and *t* cells, but the integrated signal at the soma reflects the larger dendritic field size of *n* cells, summing inputs from more and spatially more extended BCs in *n* cells. Interestingly, *d* cells had significantly larger RFs for intermediate distances between approx. 50 and 100 *μ*m than both *t* and *n* cells.

### Dendritic signals have diverse spatial and temporal response properties

To analyse temporal and spatial properties of dendritic signal integration, we used, in addition to the noise stimulus, a local (300 *μ*m diameter) and a global (≈800 *μ*m diameter) “chirp” stimulus and, because of limited recording time only for some fields, a “sine-spot” stimulus consisting of a small (60 *μ*m diameter) and medium spot (300 *μ*m) played in alternation (Fig. 2; Methods). As for the noise, the stimuli were always centred on the recording site. For the chirp stimulus, we found that in some cases, responses to the local and global chirp were almost identical (Fig. 2A, B). However, in other cases, only the local chirp stimulus resulted in an “ON” response while the global chirp resulted in an “OFF-suppressed” response, likely because of a stronger surround stimulation (Fig. 2B). In many cases, even the 60 *μ*m spot of the sine-spot stimulus was able to reliably evoke responses sometimes even stronger than the 300 *μ*m spot (Fig. 2C), indicating first, that these postsynaptic dendritic signals are sensitive to localised, small light spots, and second, that the ON component of these signals is likely dominated by excitatory inputs from only a few BCs close to the respective ROIs’ locations and activating more BCs more distant does not increase the response. We also found that the RF of different ROIs did not only vary in their spatial but also their temporal tuning, ranging from transient biphasic temporal RF to more sustained monophasic temporal RFs (Fig. 2D).

**Fig. 2.**
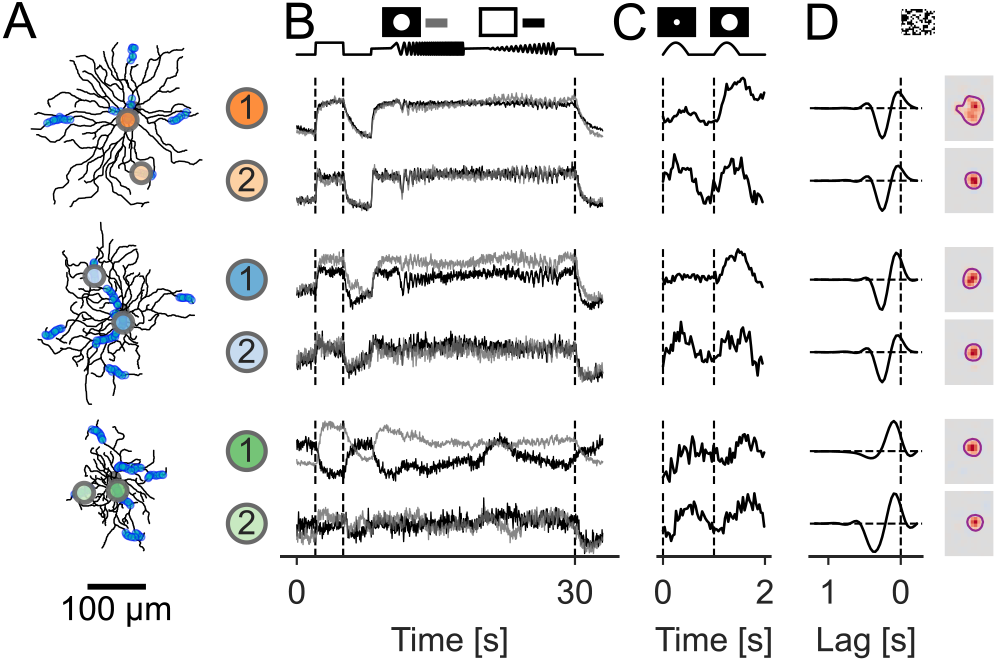
Dendritic signals have diverse spatial and temporal response properties. **A**. Cells from Fig. 1K with two example ROIs highlighted for each cell. **B-D**. Responses of example ROIs to all light stimuli used. **B**. Chirp (local: grey; global: black) response averages over repetitions. **C**. Sine-spot response averages. **D**. Dense-noise response shown as temporal (*left*) and spatial (*right*) RFs.

### sONα RGCs exhibit regional differences in dendritic signal integration

Overall, we found that the dendritic Ca^2+^ signals in response to the chirp stimuli were quite diverse, especially for the global chirp where some of the ROIs showed “OFF-suppressed” responses (Fig. 2B and Fig. S3A). This dendritic signal diversity could arise from differences in local synaptic inputs and/or electrical anatomy at the ROI (e.g. the distance to the soma). To distinguish between these possibilities, we summarised the most prominent signal features by clustering the ROIs based on their local and global chirp using a hierarchical clustering algorithm (Methods). Based on the dendrogram, we selected a distance threshold to strike a balance between simplifying the data and not merging very different responses, resulting in three clusters (Fig. S3A). The three clusters mostly differed in their transience and, interestingly, their strength of the surround suppression, and could be described as “ON-sustained” (C1), “ON-weakly-transient” (C2) and “local-ON-global-OFF-suppressed” (C3) (Fig. 3).

**Fig. 3.**
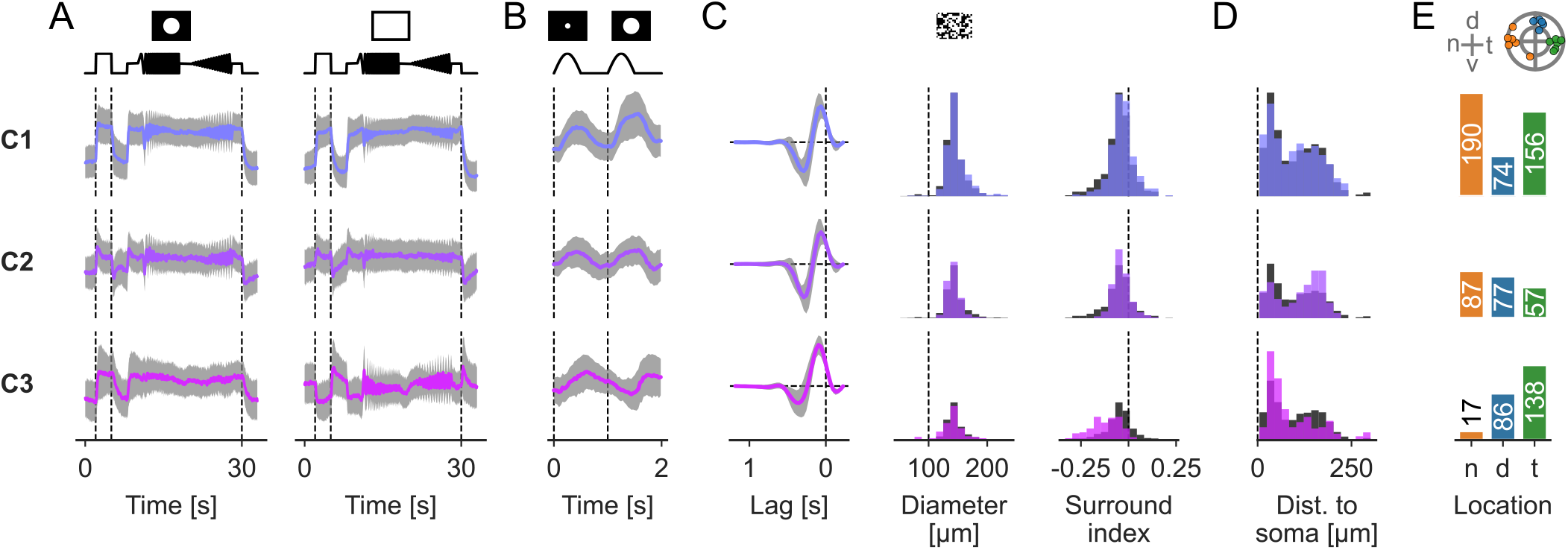
Temporal sONα show postsynaptic dendritic signals with strong surround suppression. Clustered postsynaptic Ca^2+^signals of all ROIs exceeding the quality threshold. Responses were clustered based on the local and global chirp responses. All traces used for clustering and a respective dendrogram are shown in Fig. S3A. Data is split by the three Ca^2+^clusters C1 (*top*), C2 (*middle*) and C3 (*bottom*). **A**. Local (*left*) and global (*right*) chirp responses. **B**. As in A, but for sine-spot responses. **C**. RF properties: temporal RF (*left*), spatial RF diameter (*middle*), and surround index (*right*). **D**. ROI distribution of distance to soma. **E**. ROI counts per regional group. *Top*: Cell locations for reference. **A-E**. All traces are shown as cluster mean *±* one standard deviation. For all histograms, the distribution across all clusters, scaled to cover the same area, is shown in the background (grey).

The ON-sustained cluster C1 (327 ROIs; Fig. 3, *top row*) showed a highly sustained response and only very weak surround suppression. The RF diameters of this cluster ranged from the smallest (≤ 100 *μ*m) to the largest values we observed (*>* 250 *μ*m). The responses of the ON-weakly-transient cluster C2 (268 ROIs; Fig. 3, *middle row*) also showed little surround suppression and a strong sustained component, but it was more transient than C1. Compared to the other clusters, it was found more often in distal dendrites. The local-ON-global-OFF-suppressed cluster C3 (236 ROIs; Fig. 3, bottom row) only showed a sustained ON response to the local chirp, while for the global chirp, the response was OFF-suppressed, indicating a strong surround suppression (Fig. 3A) that was also visible in the sine-spot (Fig. 3B) and dense noise responses (Fig. 3C). Many ROIs of this cluster were located at an intermediate distance to the soma (59% of C3 ROIs between 25 μm and 75 μm; C2: 32%; C1 43%) and only a few very close to it (5% of C3 ROIs closer than 25 *μ*m; C2: 10%; C3: 12%).

Next, we analysed the contribution of cells and their retinal regions to the three clusters (Fig. 3E). We found that ROIs of *n* cells were almost exclusively found in clusters C1 (65%) and C2 (30%), whereas ROIs of *d* cells were relatively evenly distributed (C1: 31%; C2: 32%; C3: 36%) and ROIs of *t* cells where mostly found in C1 (44%) and C3 (39%). Taken together this suggests a difference in surround suppression in dendritic signals between nasal, dorsal and temporal circuits.

### Nasal and temporal sONα RGCs receive different excitatory synaptic inputs

To investigate the origin of the postsynaptic surround suppression that was present in temporal (and dorsal), but not in nasal cells, we conducted a second set of experiments where we measured excitatory synaptic inputs onto the dendrites of sONα RGCs using the glutamate biosensor iGluSnFR (see Methods) and repeated the analysis from above for this dataset. Here, we restricted the cell locations to nasal (*n*) and temporal (*t*) for simplicity. We recorded signals across the dendrites (Fig. 4A) in response to the local and global chirp (Fig. 4B), the sine-spot stimulus (Fig. 4C), and the dense noise (Fig. 4D). As for the Ca^2+^ data, the RF centres were always close to the located recordings sites (distance from ROI to RF centre 30 ± 14 *μ*m; mean ± one standard deviation) (Fig. 4E). To compare the excitatory synaptic inputs of *n* and *t* sONα RGCs, we first looked at the RF sizes in relation to the ROIs’ distances to soma for both groups (Fig. 4F). We found that, close to the soma, RF sizes of nasal cells were slightly larger, while for larger distances there was no significant difference between the two groups. Next, we quantified the strength of the antagonistic surround, measured as *surround index* (see Methods), in the RFs (Fig. 4G). The RF surround was stronger in *t* cells compared to *n* cells both close to the soma and for intermediate distances, suggesting that the surround suppression observed in *t* RGCs on the postsynaptic level (Fig. 3) may originate in glutamate synaptic inputs with stronger surround suppression. To further compare the Ca^2+^ and glutamate signals, we clustered the glutamate signals using the same method as for the Ca^2+^, i.e. by clustering their local and global chirp responses (Fig. S3B). Again, we found three clusters, one ON-sustained (G1) and two local-ON-global-OFF-suppressed clusters with weak (G2) and strong (G3) suppression (Fig. 4H-L).

**Fig. 4.**
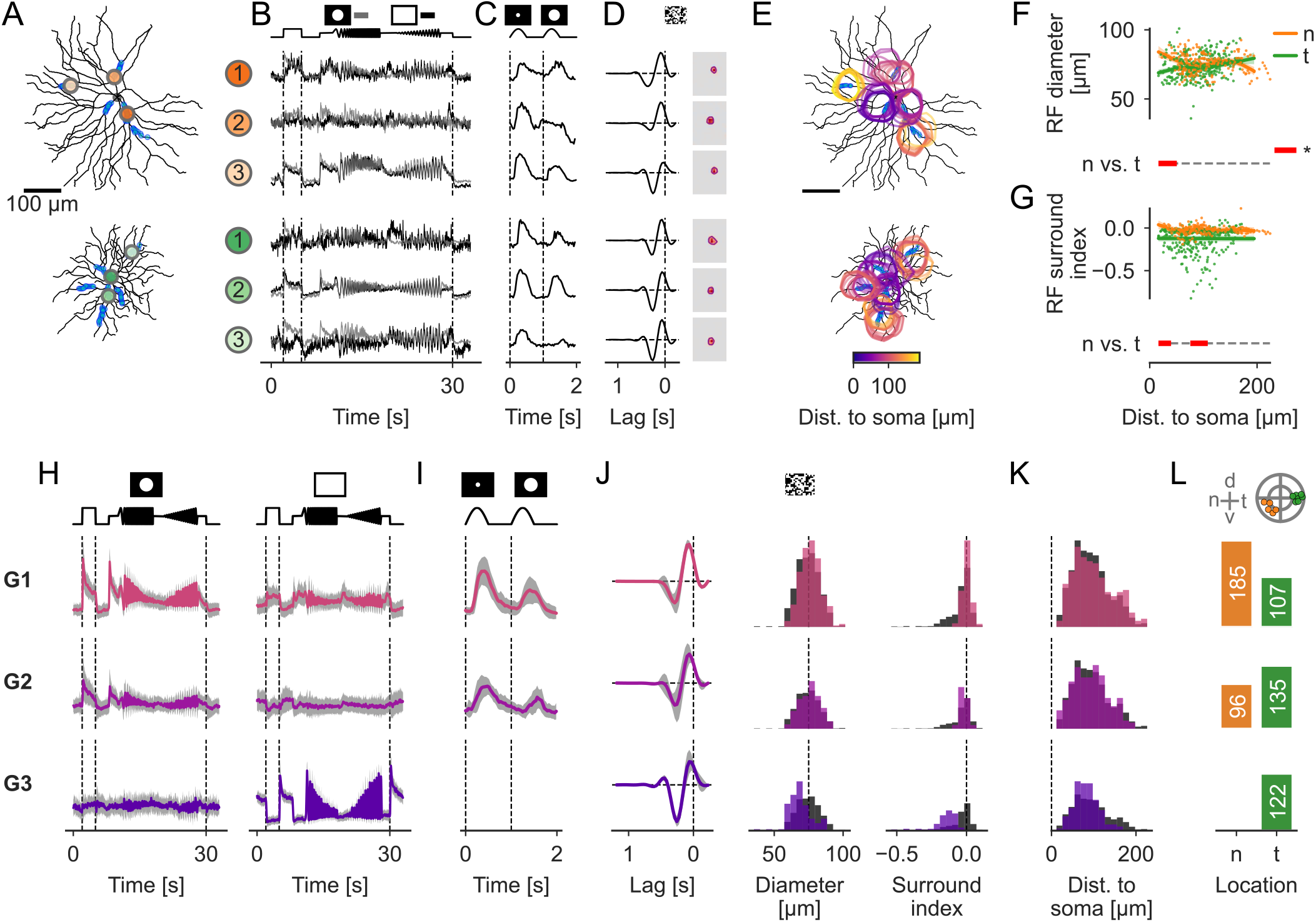
Strong surround suppression originates presynaptically in temporal sONα cells. **A**. Cell morphologies (black) with three example ROIs highlighted for each cell. **B-D**. Responses of example ROIs to all light stimuli used. **B**. Chirp (local: grey; global: black) response averages over repetitions. **C**. Sine-spot response averages. **C**. Dense-noise responses are shown as temporal (*left*) and spatial (*right*) RFs. **E**. Spatial RF outlines mapped on the morphologies (outlines colour code ROI distance to soma). **F**. RF diameter as a function of ROI distance to soma for nasal (*n*, orange) and temporal (*t*, green) cells, and fits from a Generalised Additive Model (see Methods). Distances of significant difference between *n* and *t* are highlighted (bottom; red). **G**. As in F, but for the RF surround index (see Methods). **H-L**. As in Fig. 3, but for the clustered presynaptic glutamate signals of all ROIs exceeding the quality threshold. Responses were clustered based on the local and global chirp responses. All traces used for clustering and a respective dendrogram are shown in Fig. S3B. Data is split by the three glutamate clusters G1 (*top*), G2 (*middle*) and G3 (*bottom*). **H**. Local (*left*) and global (*right*) chirp responses. **I**. Sine-spot responses. **J**. RF properties: temporal RF (*left*), spatial RF diameter (*middle*), and surround surround index (*right*). **K**. ROI distribution of distance to soma. **L**. ROI counts per regional group. *Top*: Cell locations for reference.

The ON-sustained cluster (G1; Fig. 4H-L, *top row*) had sustained ON responses for both chirp stimuli, with a preference for the smaller diameter. This cluster had the largest average RF size and the weakest RF surround suppression. The distance-to-soma distribution was relatively even, with little difference from the other clusters. The local-ON-global-OFF-suppressed cluster with weak suppression (G2; Fig. 4H-L, *middle row*) had a strong and sustained ON response for the local chirp. The global chirp response was OFF-suppressed and more variable. The preference for the smaller spot was more prominent compared to cluster G1. RF sizes were slightly smaller and surround suppression slightly stronger than for G1. The local-ON-global-OFF-suppressed cluster with strong suppression (G3; Fig. 4H-L, *bottom row*) had a weak ON response for the local chirp. The response was completely suppressed by the global chirp. In G3, RF sizes were the smallest and RF surround suppression was the strongest among the three clusters.

Taken together, similar to the Ca^2+^ clusters (Fig. 3), we found that the glutamate clusters (Fig. 4H-L) with stronger surround suppression were more frequently found in *t* cells, with ROIs from cluster G3 with the strongest suppression exclusively in *t* cells. This indicates that the stronger surround suppression observed in the dendritic Ca^2+^ signals of temporal sONα cells is at least partially inherited from the BCs inputs, likely reflecting presynaptic suppression.

### Regional adaptations in temporal sONα RGCs are well suited for prey capture

The results so far showed that sONα RGCs feature distinct regional adaptations not only in morphology but also in postsynaptic dendritic signal processing and presynaptic inputs. This was most striking in the strong surround suppression we observed in sONα cells in the temporal periphery of the retina. Notably, this region coincides with the binocular area of the mouse’s visual field, which also has been proposed to play a critical role in visually guided hunting (24). Therefore, we hypothesised that the regional adaptions in dendritic input and signal processing of sONα RGCs are an adaptation beneficial for tasks like prey capture.

To test this, we created an encoder-decoder paradigm (Fig. 5; Methods), with different sONα population models encoding scenes of a visually-guided cricket hunt reconstructed by Holmgren et al. (25), and a decoder trained to detect the presence or absence of a cricket (Fig. 5A). This encoder-decoder paradigm allowed us to compare how suited different populations of sONα RGCs are to encode the presence of a cricket. Each encoder consisted of two populations of BCs, one with weak (*w*) and one with strong surround suppression (*s*) with parameters based on the glutamate clusters G1 and G3, respectively (see Methods) and an RGC population (Fig. 5B). For simplicity, we omitted cluster G2 in the models. The two BC populations were modelled as a square grid of BCs, each with a spatial RF (2d convolution of the input), a non-linearity (generalised sigmoid) and additive Gaussian noise (Fig. 5B, C). The RGC population was modelled as a square grid of RGCs, with dendritic arbours (2D convolutions of the input) and distances dependent on the RGC population (Fig. 5B, C). The decoder was a simple convolution neural network that we trained for each encoder independently.

**Fig. 5.**
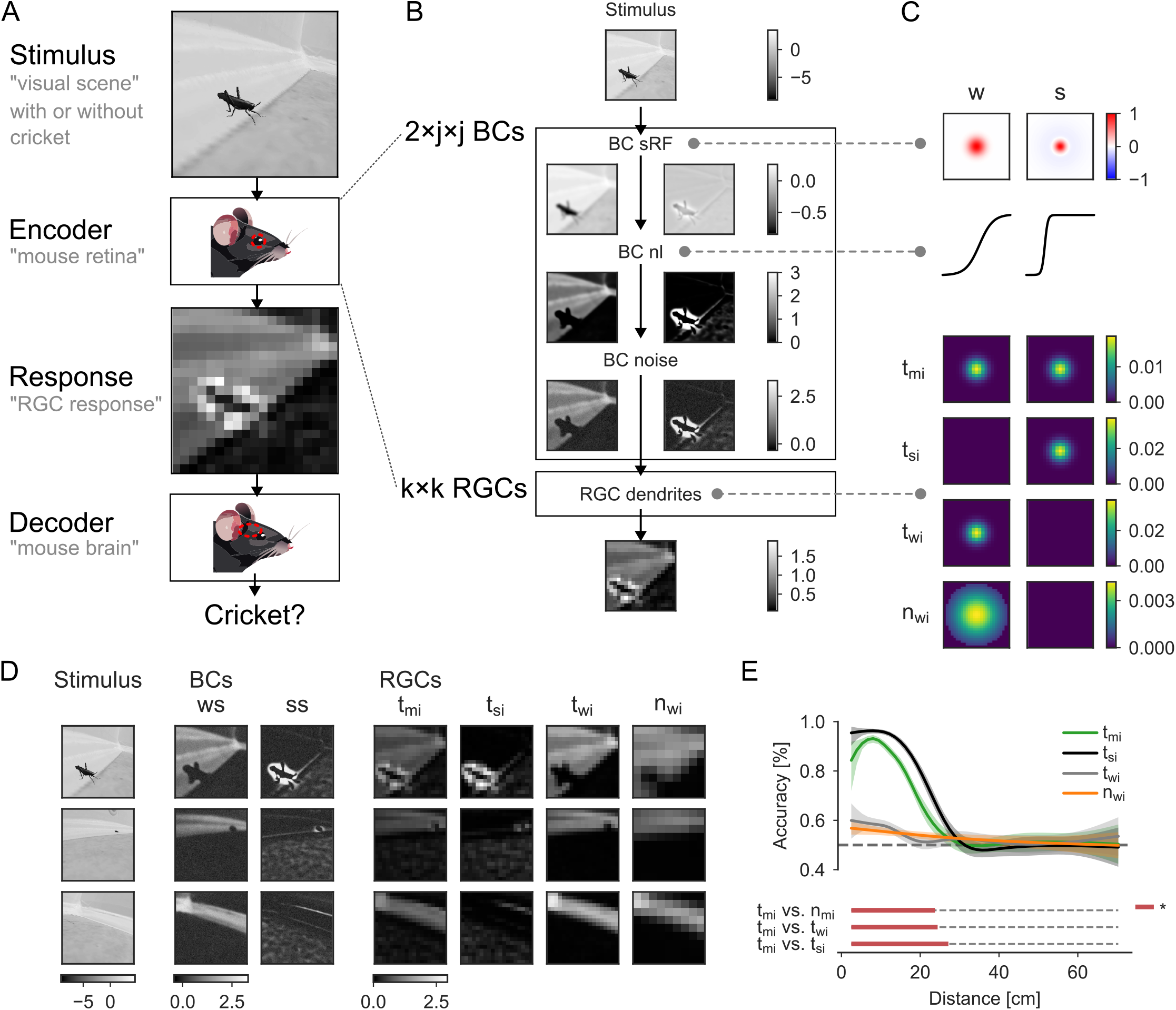
Model implementation. **A**. Encoder-decoder framework: A visual stimulus is presented to an encoder that simulates the response of an RGC population which is fed to a decoder that has to estimate if the input contains a cricket or not. **B**. Encoder structure and intermediate layer evaluations: The encoder, implemented as an artificial n euronal n etwork, i s m odelled a s t wo p opulations of *j × j* BCs, and a population of *k × k* RGCs. Each BC population consists of a 2d convolution layer (BCs’ spatial RFs), followed by a non-linearity, and a Gaussian additive noise layer. The RGC population consists of a single 3d convolutional layer (RGC dendrites), integrating the signals from the two BC populations. **C**. Encoder parameter details. *Top:* The spatial RFs and the non-linearities of the two BC populations, one with weak surround (w) and one with strong surround (s), respectively. *Bottom:* The dendritic weights of four different populations of RGCs: temporal (*t*) RGCs with inputs from either both (*t*_*mi>*_), or only from *w* BCs (*t*_*wi>*_), or *s* BCs (*t*_*si>*_); and nasal (*n*) RGCs with inputs from *w* BCs only (*t*_*wi>*_). **D**. Simulated BC and RGC responses for three example stimuli, one with a close, one with a distal and one without a cricket. **E**. Decoder accuracy fitted with a logistic GAM as a function of the cricket distance (Methods), for the four RGC populations, each tested against the *t*_*mi*_population.

We created different encoder models, based on the functional data described above and anatomical data from Schwartz et al. (26) and Bleckert et al. (22) (see Methods). The temporal RGC population (*t*_*mi>*_) we modelled received mixed inputs from both BC populations (Fig. 5C) as observed (see Fig. 4L). For this population, the cricket was clearly visible in the population encoding, especially if the cricket was very close (Fig. 5D), which is reflected in the high decoder performance for crickets closer than 20 cm (Fig. 5E). To test the role of the inhibitory surround, we compared the *t*_*mi>*_ RGC population to a population receiving only strong surround (*t*_*si>*_) or only weak surround input (*t*_*wi>*_) (Fig. 5C-E). The decoder performed best for the temporal population with strong surround inputs only with significantly higher accuracies than the model with mixed inputs (Fig. 5E), with high performance for close cricket and a sharp decay in accuracy at around ≈ 20 cm, similar to the mixed inputs model. Without inputs from BCs with strong surround, the decoder performance was significantly worse (Fig. 5E).

Finally, we compared the temporal RGCs to a model of nasal RGCs, with larger dendritic arbours, i.e. pooling inputs from more BCs, larger distances between RGCs and with inputs from BCs with weak surround only (see Fig. 4L). For this population, the decoder performance was very similar to the performance of the temporal population with weak surround only, indicating only a minor effect of dendritic size and spacing on the observed cricket detection performance.

Taken together, our results suggest that signal integration at the level of temporal sONα dendrites together with changes in the presynaptic circuits indeed are tuned for detecting small objects such as moving insects and, hence, could improve visual hunting performance in mice.

## Discussion

In this study, we investigated the function of the previously reported higher density of sONα cells in the temporal mouse retina (22). Specifically, we asked if this anatomical adaptation goes beyond higher spatial sampling (i.e. higher cell density and smaller dendritic arbours) and is accompanied by distinct functional changes. To this end, we looked into the dendritic signal processing of sONα RGCs in different regions of the animal’s retina.

We found that dendritic Ca^2+^ signals in nasal sONα RGCs were mostly ON sustained with modest surround suppression, as it was reported for this cell type in earlier studies (e.g. 21, 26, 27). In contrast, temporal sONα RGCs additionally exhibited dendritic Ca^2+^ signals with strong surround suppression in more than a third of the dendritic segments we measured from. This strong surround suppression was already present in the excitatory synaptic inputs onto these RGCs, pointing to the involvement of presynaptic mechanisms.

Using computational population models of these cells, we analysed how nasal and temporal sONα RGCs encode movies of small moving objects such as crickets. Our modelling results indicate that the observed differences in synaptic inputs could be a regional adaptation beneficial for tasks like visually guided hunting.

### Regional adaptations in the retina

Regional cell-type specific adaptations (reviewed in 1, 28) have long been studied in the retina. For example Boycott and Wässle (29) observed in cat that some RGC types become denser while significantly decreasing in dendritic field size towards the central retina, with the highest densities and smallest dendritic arbours in the *area centralis*. Such regional changes in cell density/dendritic arbour size are common in vertebrates and typically linked to visual acuity – the denser the mosaic of a cell type, the higher its spatial resolution. Regional adaptions occur also upstream of the RGCs; for instance, cone photoreceptors (cones) in the primate fovea are slower than peripheral cones and hence, shape foveal perception (30).

In mice, it has been observed that a large fraction of eye movements are compensatory and counteract their head/body motion (31). Therefore, it is expected that mice stabilise the visual scene on their retina with respect to the cardinal axes of the world. As a result, prominent scene features, such as the horizon, tend to fall on distinct parts of the mouse retina, which, in turn, enables its partitioning into specialised regions. This is different from, for example, primates, which also feature a specialised region – the fovea – but use it to “scan” the visual world (32, 33). It has therefore been proposed that the ventral mouse retina, which is “looking up” and covers the upper visual field, may be specifically tuned for detecting birds of prey in the sky (reviewed in 1, 28).

Recent mouse studies have revealed several regional adaptations at all retinal levels, including a prominent opsin expression gradient along the dorsal-ventral axis (7–10), region-dependent axonal territory sizes in OFF bipolar cells (34), and an overall lower RGC density in the dorsal retina (16, 17). There were also several RGC type-specific regional adaptations reported, such as distinct density distributions (17, 19, 22, 35), at times associated by changes in morphology (18, 22, 34). Adaptations on the functional level have only been reported for a few RGC types so far, for instance, for the transient OFF alpha cells (tOFFα; 21) and the JAM-B cells (18), both of which vary in their response along the dorso-ventral axis. tOFFα cells were reported to feature more sustained light responses in the dorsal vs. the ventral retina (36), while JAM-B cells change from being (modestly) direction-selective in the dorsal to colour-opponent in the ventral retina (37, 38). In the latter, the functional change is accompanied by a change in dendritic arbour morphology (18). In the present study, we have identified another RGC type, the sONα, that exhibits fine-grained regional functional differences – supporting the view that local adaptations of functional properties and, hence, distinct roles in different regions of the retina, are not the exception but the rule in animals like mice. Such functional regionalisation further adds to the already astonishing diversity of RGC signals (27, 39) in the mouse retina.

### Functional properties of sONα cells

The sONα RGC can be distinguished relatively easily from other RGC types based on their highly sustained somatic ON responses, large soma sizes, and SMI-32 immunoreactivity (20, 21, 23, 40). For this reason, they are ideally suited to investigate regional adaptations of retinal circuits.

sONα RGCs have a high base firing rate under steady illumination (20, 21, 41) driven by excitatory synaptic inputs originating mostly in type 6 and type 7 BCs (26, 42, 43). While sONα cells express low levels of melanopsin and are, hence, intrinsically photosensitive (44), their light response is dominated by the synaptic inputs (40). The surround receptive field of sONα is antagonistic (20, 40), presumably by suppression of excitatory presynaptic inputs (20, 43).

Previous studies have already shown the systematic variation in sONα RGCs morphology (22, 23), with a temporal “hotspot” where the cells have the highest density and smallest dendritic fields. Bleckert et al. (22) also showed that the temporal cells have a higher coverage factor than nasal sONα cells, potentially increasing the spatial acuity of the cell population.

In this study, we found that nasal and temporal sONα cells exhibit distinct dendritic signal processing, most prominently visible in the surround strengths of their dendritic glutamatergic inputs.

Very recently, Hsiang et al. (45) have shown that type 7 BCs have a very strong surround, with the centre and surround responses virtually cancelling each other for spots (bright; flashed for 1.5 s) of a diameter between 200 *μ*m and 400 *μ*m and completely sign-inverted responses for spots larger than 600 *μ*m, consistent with our glutamate clusters G2 and G3 (Fig. 3H, I). For type 6 BCs, they found that the strongest response is achieved for spot sizes of around 100 *μ*m, while spots larger than 400 *μ*m resulted in substantially weaker ON responses to the light increment and a below baseline suppression for the light decrement, likely corresponding to our glutamate cluster G1 (Fig. 3H, I). Hence, our results suggest that temporal sONα RGCs receive more synaptic input from type 7 vs. type 6 BCs compared to nasal sONα cells.

In another recent study, Swygart et al. (43) performed similar experiments to ours, where they also recorded from sONα cells in different regions of the retina. In contrast to our results, they did not find a significant difference in surround suppression between different locations of the retina. Several factors may explain these seemingly inconsistent findings. First, the recordings, for which Swygart et al. reported the retinal regions, were performed under scotopic conditions, while we recorded at low photopic light levels. Second, Swygart et al. used exclusively a blue LED (450 nm), which drives mostly rods and M-cones, whereas we used a UV (390 nm) and a green (575 nm) LED to stimulate both S-and Mcones, respectively. A colour-dependency of the surround strength seems plausible given the diverse and location-dependent colour preference of the sONα cell’s surround reported along the dorsal-ventral axis (14). Finally, Swygart et al. (43) electrically recorded from sONα somata, whereas we recorded directly from dendrites. It is possible, that the surround suppression we observed on dendritic segments becomes less pronounced in the somatic signal due to the integration of all the dendritic signals. If so, dendritic surround suppression would play a different role as a canonical antagonistic surround. One such role may be to render the RF’s spatial structure less linear (26), which could make the cells more sensitive to spatial contrasts.

Alternatively, the dendritic surround suppression could also be a means to decrease the overall excitation of temporal sONα cells for very bright light levels. Berry et al. (46) showed that, under photopic conditions and full-field stimulation, intrinsically photosensitive RGCs, including clusters that likely correspond to the sONα, encode visual information relatively poorly compared to mesopic conditions. Especially, if the surround is mostly activated during higher light levels, the suppressed surround could counteract over-excitation and stabilise the response of temporal sONα for different light levels.

The results reported by Swygart et al. (43) may also provide another mechanism for the surround suppression in synaptic inputs and postsynaptic Ca^2+^ signals we observed. Their data suggest that type 6 BCs have multiplexed outputs, with ribbon synapses featuring weak and strong surround suppression, mediated by amacrine cells, in individual BCs. While they use this to explain the stronger surround suppression in PixON (EyeWire: 9n) RGCs (47) compared to sONα RGCs, this may also be the mechanism that enables increasing dendritic surround suppression in temporal sONα cells with minimal changes in the circuit. Temporal sONα cells would not even have to adjust the ratio of their presynaptic partners to receive stronger surround suppression inputs but only connect to different ribbons of the same type 6 BCs.

### A role for sONα RGCs in visually-guided hunting

Several studies have shown that mice use their vision to hunt prey (24, 25, 48–50). Especially the binocular region in the temporal mouse retina – where sONα cells have their highest density – seems to play a critical role (24, 25). Johnson et al. (24) showed that of the 40+ RGC types in mice, only a subset of nine types makes ipsilateral connections to the brain. Moreover, the authors showed that from these nine types only five, including the sONα RGC, have reliable responses to a stimulus mimicking a moving insect, suggesting that these RGC types are critical for successful hunting. In a related study, Holmgren et al. (25) showed that mice bring the image of their prey on a relatively small spot in the temporal retina with high accuracy, coinciding with the region where sONα RGCs have the highest density. Together, this suggests that sONα RGCs in this temporal high-density region play a role in hunting and that any regional functional adaptations relate to this and related behavioural tasks. In our study, we provide further evidence that temporal sONα RGCs play a dedicated role in visually-guided prey capture, by demonstrating that the regional adaptions we found in these cells can indeed be advantageous for such a task.

It should be noted, though, that we do not propose that hunting behaviour is supported by a single RGC type; certainly, other RGCs also play vital roles in such a complex behavioural task. For example, OFF RGCs in particular would be well suited to complement the signals from sONα RGCs. Among the nine ipsi-laterally projecting RGCs, there are only two OFF types: sOFFα (EyeWire: 1wt) and tOFF (EyeWire: 4i) (24); with the sOFFα being especially interesting because its highest density region coincides with that of sONα (22).

Beyond the retina, Krizan et al. (51) recently showed that narrow-field neurons in the superior colliculus of mice are used for hunting and that these neurons receive input from both direction and non-direction selective RGCs. Interestingly, eliminating retinal direction selectivity did not affect the animals’ hunting behaviour and success, suggesting that non-direction selective RGCs, therefore potentially sONα RGCs which also project to the superior colliculus (44, 52, 53), provide critical input for prey capture to these neurons.

### Linking the retina to behaviour

In this study, we used an encoder-decoder paradigm that allowed us to analyse the effects of different inputs and dendritic field sizes of sONα RGCs systematically and efficiently. The assumptions of the encoder, i.e. the sONα population model, were rather conservative and reduced to the main regional differences we observed, namely the morphological differences and the different levels of surround suppression in the presynaptic input.

For the sake of simplicity, we excluded other factors that differ between the populations, such as temporal effects, inhomogeneities (e.g., non-Gaussian) in the spatial structure (26) of the RFs, and other RGC types. Therefore, our encoder-decoder model may well underestimate the performance of the biological counterpart. In a future study, it would be interesting to see how the model’s performance in the encoding-decoding task changes when including the signals of any of the aforementioned OFF types. Another limitation of our model may come from the videos we used. While they were reconstructed from freely moving mice hunting crickets, and thus provided an accurate visual input as seen by these mice, the videos were recorded in an artificial, well-lit environment. Hence, the visual scene statics, in particular, concerning background and illumination (“backdrop”, see 54), were far from naturalistic.

In general, it is difficult to relate the retinal output to something as complex as behaviour. To address this problem, there are several approaches in the literature that can be broadly grouped as follows. In some studies, the researchers looked at behavioural data and tried to link their observations to previous findings about the retina (e.g., 25, 55). Other studies used functional recordings in response to either artificial or natural stimuli and tried to draw conclusions about the behavioural relevance (e.g., 19, 56). The aforementioned study by Johnson et al. (24) is particularly interesting in this respect because they used both behavioural data and functional retinal recordings to find the most important RGC types for this task. With the encoder-decoder paradigm used in this study, we were able to make another link from functional retinal recordings to hunting behaviour. We believe that this approach of combining natural scenes, a retinal encoder model, and a decoder trained on a simple task with direct behavioural relevance, offers yet another angle to address the crucial question of functional relevance.

## Methods

### Animals and preparation

Mice used in this study were purchased from Jackson Laboratory and housed under a standard 12 h day/night cycle with 22°C, 55% humidity. Mice aged 8 to 12 weeks of either sex were used for all experiments. For the Ca^2+^ recordings, we used the wildtype line (C57BL/6J, JAX 000664; n=12 animals). For the glutamate recordings, we used the crossed *B6;129S6-Chattm2(cre)Lowl/J* (ChAT:Cre, JAX 006410) × B6.Cg-*Gt(ROSA)26Sortm9(CAG-tdTomato)Hze/J* (Ai9tdTomato, JAX 07909) mouse line that we virus-injected intravitreally to express iGluSnFR in the retina (n=5 animals; see virus injection). All animal procedures were approved by the governmental review board (Protocol numbers: CIN 3/18 G, animal protocol from 31.10.2016, Regierungspräsidium Tübingen, Baden-Württemberg, Konrad-Adenauer-Str. 20, 72072 Tübingen, Germany) and performed according to the laws governing animal experimentation issued by the German Government.

Mice were dark-adapted ≥ 2 h before tissue preparation, then anaesthetised with isoflurane (Baxter, Hechingen Germany), and killed with cervical dislocation. We marked the dorsal side of each eye with dye before quickly enucleating them in carboxygenated (95% O_2_, 5% CO_2_) artificial cerebral spinal fluid (ACSF) solution containing (in mM): 125 NaCl, 2.5 KCl, 2 CaCl_2_, 1 MgCl_2_, 1.25 NaH_2_PO_4_, 26 NaHCO_3_, 20 glucose, and 0.5 L-glutamine (pH 7.4). After removing the cornea, sclera and vitreous body, the retina was flattened on an Anodisc (0.2 *μ*m pore size, GE Healthcare, Pittsburgh, PA) with the ganglion cell side facing up and then transferred to the recording chamber of the microscope, where it was continuously perfused with carboxygenated ACSF (at 35°C and 4 ml min^*−>*^1). All experimental procedures were carried out under very dim red light.

### Virus injection

Before virus injection, mice (5-7 weeks) were anaesthetised with 10% ketamine (Bela-Pharm GmbH, Germany) and 2% xylazine (Rompun, Bayer Vital GmbH, Germany) in 0.9% NaCl (Fresenius, Germany). One *μ*l of AAV9.hSyn.iGluSnFR.WPRE.SV40 (Penn Vector Core, PA, USA) was loaded into a Hamilton syringe (syringe: 7634-01, needle:207434, point style 3, length 51 mm, Hamilton Messtechnik GmbH). Then, the syringe was fixed on a micromanipulator (M3301, World Precision Instruments, Germany) and the virus was slowly (1 *μ*l/5 min) injected into the vitreous body. Virus-injected mice were used for recordings after 3 weeks.

### Single-cell micro-injection

To visualise blood vessels and avoid them when filling individual RGCs, 5 *μ*l of a 50 mM sulforhodamine-101 (SR-101, Invitrogen/Thermo Fisher Scientific, Dreieich, Germany) stock solution was added per litre ACSF solution. Sharp electrodes for single-cell injection were pulled on a P-1000 micropipette puller (Sutter Instruments, Novato, CA) with resistances ranging between 70 and 130 M!1. For Ca^2+^ indicator loading, Oregon Green BAPTA-1 (OGB-1, hexapotassium salt; Life Technologies, Darmstadt, Germany; 15 mM in water), a synthetic Ca^2+^ indicator dye with high Ca^2+^ affinity (K_D_=170 nM; Invitrogen) and comparatively fast kinetics (57), was loaded into individual RGCs using the single-pulse function (500 ms, -10 nA) of a MultiClamp 900A amplifier (Axon Instruments/Molecular Devices, Wokingham, UK). For the visualisation of single RGC morphologies, while recording iGluSnFR signals, 10 mM of Alexa Fluor 594 (Invitrogen/Thermo Fisher Scientific, Dreieich, Germany) was injected into individual RGCs using the same method for Ca^2+^ indicator loading. To allow the cells to fill and recover, we started recordings 1 h post injection.

### Two-photon imaging and light stimulation

We used a MOM-type two-photon microscope (designed by W. Denk, MPI, Martinsried; purchased from Sutter Instruments/Science Products) as described previously (58). Briefly, the system was equipped with a mode-locked Ti:Sapphire laser (MaiTai-HP DeepSee, Newport SpectraPhysics, Darmstadt, Germany), green and red fluorescence detection channels for OGB-1 (HQ 510/84, AHF, Tübingen, Germany) and SR-101/tdTomato (HQ 630/60, AHF), and a water immersion objective (W PlanApochromat 20×/1,0 DIC M27, Zeiss, Oberkochen, Germany). For all scans, we tuned the laser to 927 nm, and used custom-made software (ScanM, by M. Müller, MPI, Martinsried, and T.E.) running under Igor Pro 6.3 for Windows (RRID:SCR_000325; Wavemetrics, Portland, OR). Dendritic Ca^2+^ and glutamate signals were recorded with 64 × 16 pixel image sequences at 31.25 Hz. We acquired high-resolution mythology stacks using 512 × 512 pixel image stacks with 0.8 or 1.0 *μ*m z steps.

For the light stimulation, a digital light processing (DLP) projector (lightcrafter, DPM-E4500UVBGMKII, EKB Technologies Ltd) was used to display visual stimuli through the objective onto the retina, whereby the stimulus was focused on the photoreceptor layer (58, 59). The lightcrafter was equipped with a light-guide port to couple in external, band-pass filtered green and UV light-emitting diode (LEDs; green: 576 BP 10, F37-576; UV: 387 BP 11, F39-387; both AHF/Chroma). The band-pass filter was used to optimise the spectral separation of mouse M- and Sopsins (390/576 Dualband, F59-003, AHF/Chroma). The LEDs were synchronised with the scan retracing of the microscope and intensity-calibrated to range from approx. 0.1 × 10^3^ (black background) to 20.0 × 10^3^ (white full field) photoiso-merisations (P*) s^−1^ cone^−1^.

The light stimulus was carefully centred before every experiment, ensuring that its centre corresponded to the centre of the microscope’s scan field. For all experiments, the tissue was kept at a constant mean stimulator intensity level for ≥ 15 s after the laser scanning started and before light stimuli were presented. Light stimuli were generated and presented using the Python-based software package QDSpy (RRID:SCR_016985). We used four types of light stimuli:

1. Binary dense noise (20 × 15 matrix of 30 *μ*m per pixel; each pixel displayed an independent, balanced random sequence at 5 Hz for 5 min) for spatio-temporal receptive field (RF) mapping. The pixel size was chosen to be slightly smaller than the RF centre of single BCs (38–68 *μ*m in diameter; 60), allowing to estimate RGC dendritic RFs at single-BC resolution.
2. Full-field (800 × 600 *μ*m) chirp, consisting of a bright step and two sinusoidal intensity modulations, one with increasing frequency (0.5–8 Hz) and one with increasing contrast.
3. Local chirp; like (2) but as 300 *μ*m-diameter spot.
4. Sine-spot; a sequence of light spots, 60 *μ*m and 300 *μ*m in diameter, with the intensity following a clipped sine wave (max(0, *A* sin(*π t*)), where A is the maximum intensity) resulting in 1 s of light followed by a 1 spause before the next spot.

### Reconstruction of cell morphologies and retinal location

Immediately following the recording, we captured the full dendritic structure of the RGC using a high-resolution image stack. Through semi-automatic neurite tracing techniques, we reconstructed the cell skeletons of the documented RGCs. If the imaged region was not flat, we adjusted the image stack and outlined the cell (27). All subsequent analyses, including the retrieval of morphological parameters (detailed below), were conducted using custom Python scripts.

After the functional and morphological recordings, we recorded the outline of each retina. To reconstruct the positions of cells within the retina from these recordings, we used `Retistruct` (61). In a few cases, retinal outlines were incomplete, e.g. a wing was missing, and manually adjusted. To define the orientation of the retina, we marked the dorsal side of the eye before enucleation and used this mark to make a dorsal cut in the retina towards the optic disc. In `Retistruct`, this cut was then set to be dorsal. Furthermore, to better align our data with previously published data (i.e. 17, 22), we corrected for the mean angular displacement of 22.1° between defining dorsal based on dorsal marks vs. defining dorsal based on the nasal choroid fissure (e.g., 62).

Recorded dendrites and the respective ROIs (see below) were not necessarily well-aligned with the cell morphology reconstructed later. Hence, we aligned each recording field with the respective morphology as follows: We averaged the recording field over time, normalised this average to be between zero and one (clipping values smaller than the 20^th^ and larger than the 90^th^ percentile), and rotated it to match the orientation (with the angle taken from the MOM setup) in the reconstructed skeletons. Next, we cropped the skeleton to a region of ≈ 250 × 250 *μ*m around the expected position (taken from the raw MOM-setup position readouts) of the recording field, blurred it using a Gaussian 3D filter, and normalised the result to range between zero and one. We iterated over the z-layers of the crop and used the `matchTemplate` function of `opencv-python` to evaluate how well the field matched with the crop – measured as the Mean Squared Error (MSE) – for all possible xy-positions within each layer. To penalise matches far away from the expected position, we added an Euclidean distance term to the MSE loss. Finally, we used the xyz-position with the lowest loss as the field’s position with respect to the morphology. ROIs within the field were then projected to the closest dendritic branch based on their Euclidean distance to the non-blurred skeleton (63).

### Immunohistochemistry and confocal microscopy

After single-cell recordings, the retina was removed from the anodisc and mounted to a new filter paper (0.8 *μ*m pore size, Milipore). Then the retina was fixed using 4% paraformaldehyde in 0.1 M phosphate-buffered saline (PBS) for 20 minutes at 4°C, washed with 0.1 M PBS (6×20 minutes at 4°C) and blocked with blocking solution (10% normal goat serum (NGS) and 0.3% Triton X-100 in 0.1 M PBS) overnight at 4°C. Afterwards, the samples were incubated with primary antibodies (anti-SMI-32, 1:100, Biolegend, USA, #801701, and anti-RBPMS, 1:500, Phosphosolution, USA, #1832-RBPMS) in 0.3% Triton X-100 and 5% NGS in 0.1 M PBS for 3 days at 4°C. The samples were then washed with 0.1 M PBS (6×20 minutes at 4°C) and incubated with secondary antibodies conjugated to Cy3 and Alexa Fluor 488 (1:500, ThermoFisher, Germany) in 0.1 M PBS overnight at 4°C. After another washing step (6 × 20 minutes at 4°C), the retina mounted on filter paper was embedded in Vectashield (Vector Laboratories, USA) on a glass slide and covered with a coverslip. Confocal images were taken using a Leica TCS SP8 confocal microscope equipped with 488 and 552 laser lines. Images were taken with HC PL APO 20x/75 and 40x/1.3 oil objectives. Confocal image stacks were aligned with the 2P image stacks, projected to 2D using weighted z-projections, and brightness and contrast adjusted, using custom Python scripts.

### Data analysis and software environments

Image extraction and semi-automatic region of interest (ROI) placement (see below) were performed using Igor Pro 6.37. A time marker in the recorded data was used to align the visual stimulus with 2 ms-precision. All subsequent steps were performed using custom Python code built around a database implemented using DataJoint (64). The code for the database, the analysis, and the model will be made available here: https://github.com/eulerlab/s-on-alpha_dendrites. Package versions for the analysis are listed in Table 1 and for the model (see below) in Table 2.

**Table 1.** Software analysis.

| Package | Version |
| --- | --- |
| Python | 3.10.12 |
| datajoint | 0.14.1 |
| numpy | 1.26.2 |
| pandas | 2.2.1 |
| scipy | 1.11.4 |
| scikit-learn | 1.3.2 |
| pingouin | 0.5.4 |
| statsmodels | 0.14.1 |
| matplotlib | 3.8.2 |
| seaborn | 0.13.2 |
| R | 4.3.2 |
| mgcv | 1.9-0 |
| itsadug | 2.4 |

**Table 2.** Software model.

| Package | Version |
| --- | --- |
| Python | 3.11.5 |
| tensorflow | 2.15.0 |
| keras | 2.15.0 |
| R | 4.2.0 |
| mgcv | 1.9-0 |
| itsadug | 2.4 |

#### Regions of interest

For each field, ROIs were extracted based on dense noise responses as follows (63). First, we computed the standard deviation (s.d.) of the fluorescence intensity for each pixel over time, generating a s.d. image of the time-lapsed image stack. Pixels with a s.d. at least twice the mean s.d. of the field were considered dendritic pixels. Then, the time traces of the 100 dendritic pixels with the largest standard deviations were extracted and cross-correlated. Finally, we grouped neighbouring pixels (within a distance of 3 *μ*m) with *ρ* > *ρ*_Threshold_ into one ROI, where *ρ*_Threshold_ was the mean of the resulting cross-correlation coefficients (*ρ*). In the case of the iGluSnFR data, we drew a dendrite mask manually based on the dendrite in the red channel *before* we calculated the s.d. of the time-lapsed recording image.

Only pixels in the dendrite mask were used for ROI placing as described above and further analysis. For the Ca^2+^ data we also defined “field ROIs” and “proximal dendrite ROIs”: A field ROI was defined as the combination of all ROIs within a field. A field ROI was also a proximal dendrite ROI if the medium dendritic distance to the soma (see below) of all ROI within the field is smaller than 50 *μ*m. For cells with multiple proximal dendrite ROIs, we used the one resulting in the largest RF estimate.

#### Signal processing

After ROI placement, the respective Ca^2+^ or glutamate traces were extracted. For each ROI, we computed raw traces *r*_raw_ as means of all ROI pixels. These raw traces were detrended by subtracting a smoothed version of the respective trace *r*_*smooth*_, computed using a SavitzkyGolay filter (65) of 3^*rd*^ polynomial order, from the raw traces *r*_*detrend*_ = *r*_*raw*_ − *r*_*smooth*_. The window length of this filter was 10 s for the sines-spot stimulus and 60 s otherwise.

Detrended traces were then normalised by subtracting the median baseline signal before stimulus onset at time *t*_0_ and by dividing by the s.d. of the signal:

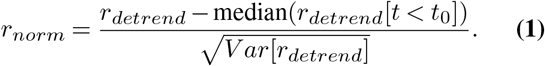

Finally, normalised responses were averaged ⟨*a* = *r*_*norm*_⟩ _*R*_over stimulus repetitions *R*.

#### Receptive field estimation

We mapped RFs of RGCs using the Python toolbox RFEst (66). The binary dense noise stimulus (20 × 15 matrix, 30 *μ*m pixels, balanced random sequence; 5 Hz) was centred on the recording field. Normalised traces *r*_*norm*_ were slightly low-pass filtered *r*_*filt*_ = *LP* (*r*_*norm*_) using a Butterworth filter (*f*_cutoff_ =3 Hz for Ca^2+^ and *f*_cutoff_ =5 Hz for glutamate), which improved the yield of high-quality RF estimates. Finally, temporal positive-only gradients were computed for each trace:

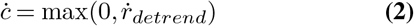

The stimulus *X*(*t*) was upsampled to the trace sampling rate of 31.25 Hz.

Spatio-temporal RFs ***F*** (*x, y, τ*) were computed from spline-based linear Gaussian models that were optimised with gradient descent to minimise the following loss:

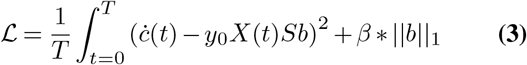

where *S* is a cubic regression spline basis, *y*_0_ is the inferred intercept, *b* are the inferred RF weights, and *β* is the weight for the L_1_-penalty on *b* to enforce sparsity in the RF. The RF was defined as ***F*** (*x, y, τ*) = *Sb. S* was defined by the number of knots in space and time (*k*_*x*_, *k*_*y*_, *k*_*t*_), corresponding to the dimensions *d* of the spatio-temporal RF (*d*_*x*_, *d*_*y*_, *d*_*t*_) = (45, 20, 15). We set to (*k*_*x*_, *k*_*y*_, *k*_*t*_) = (10, 12, 9) for Ca^2+^ and (*k*_*x*_, *k*_*y*_, *k*_*t*_) = (10, 16, 12) for glutamate. Further, we set *β* = 0.005.

Models were trained for at least 100 steps and a maximum of 2,000 steps. If the loss did not improve for 5 steps, training was stopped and the parameters resulting in the lowest loss were used as the final model.

We decomposed the RFs into a temporal *F*_*t*_(*τ*) and spatial ***F*** _*s*_(*x, y*) component using singular value decomposition and scaled them such that max(|*F*_*t*_|) = 1 and max(|***F*** _*s*_|) = max(|***F*** |). RF quality was computed as:

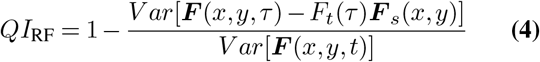

Only RFs with *QI*_RF_ > 0.35 were used for the analysis. Next, spatial RFs ***F*** _*s*_ were linearly up-sampled by a factor of 5. To estimate the RF centre outline and size, we fitted contour lines at levels 0.25, 0.3, and 0.35 using `matplotlib.pyplot.contour`. If, for all levels, there was at least one contour that covered at least 80% of the area covered by all contour lines, the largest contour line at level 0.25 was used as the RF centre outline. Other RF fits were discarded. The RF diameter was defined as the diameter of a circle covering the same area as the RF centre outline. An additional outline was drawn around this centre outline with a 20 *μ*m distance to define the inner border of the *RF surround*. The *RF surround index SI* was defined as:

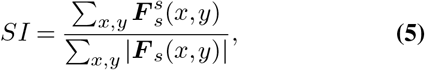

where the weights of the spatial RF surround 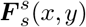 were defined as:

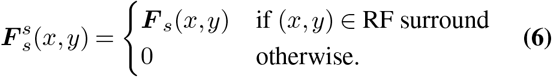

The *SI* = 1 is therefore a value between − 1 and +1 and measures the weight and sign of the surround relative to spatial RF as a whole. In some cases, the reconstructed position of the morphology was not ideally aligned with the recorded field positions. For plotting the ROI RFs on the morphology, we therefore subtracted the mean offset, i.e. the offset between field RF centre and field centre, of all field RFs per cell.

#### Quality filtering

To measure response quality for repeated stimuli (both chirps and sine-spot), we computed a signal-to-noise-ratio (SNR) quality index (27):

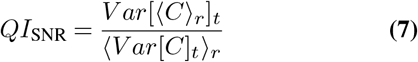

where *C* is the *T* -by-*R* response matrix (time samples by stimulus repetitions) and ⟨ ⟩_*x*_ and *V ar*[]_*x*_ denote the mean and variance across the indicated dimension *x*, respectively. We only included ROIs that had a good SNR for the local (l) or global (g) chirp:

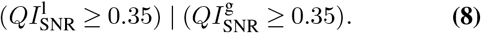

#### Functional clustering

We clustered the Ca^2+^ and glutamate datasets independently, using the same method. For both datasets, we downsampled averages from local and global chirp responses by averaging the signal over every four consecutive time points, concatenated the local and global chirp, and clustered them using Ward hierarchical clustering, implemented in `scikit-learn` (67). We used an Euclidean distance metric and a threshold of 110 that we selected based on the respective dendrograms, resulting in three clusters for both datasets.

#### Morphological metrics

Soma size was defined as the soma area in the image frame, where the soma appeared the largest. Dendritic field area was defined as the area spanned by a convex hull around the z-projected skeleton of a cell; the respective diameter was defined as the diameter of a circle with an equivalent area. The distance to soma for an ROI was defined as the length of the shortest path from an ROI to the soma centre along the dendritic arbour and computed with `MorphoPy` (68).

#### Statistical analysis

We used Generalised Additive Models (GAMs) to analyse response properties as a function of distance to soma. GAMs are an extension of Generalised Linear Models that allow linear predictors to depend on smooth functions of the underlying variables. Here we used the following GAM model:

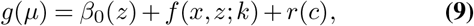

where the outcome variable *y* has expectation *μ, g* is a link function, *β*_0_ is the intercept per group *z, x* is the predictor variable, *f* is a smooth function, and *r* is a random effect per cell *c*. GAMs were implemented in R using the mgcv package. The smooths *f* were penalised cubic regression splines with dimension *k*, where lower values of *k* mean smoother fits. For each fit, we compared different values for *k*, models from the Gaussian or scaled t-distribution family, and models with and without random effect *r*. We selected the best model based on the Bayesian information criterion (BIC) and diagnostic plots. To compare the differences between groups we used `plot_diff` of the `itsadug` R package while excluding the random effect per cell. For the comparison of 2 groups we used 95% confidence intervals (CIs); for 3 groups we used 98.3% CIs, to adjust for multi-comparison.

### Model

We created population models of sONα RGCs. We derived the model parameters from our data and previously published data (see below). To simulate the differences between nasal and temporal sONα RGCs, we used different parameters for dendritic wiring and RGC spacing (see below). We used these models to encode visual scenes from freely moving mice that were hunting crickets in an arena, recorded by Holmgren et al. (25). The visual scenes we used were reconstructions of “eye views”, i.e. projections of the visual scene onto the retina, including the body, head, and eye movement of the mouse.

#### Data

The eye views were generated similarly to Holmgren et al. (25), but with 67 frames per second. To minimise projection artefacts, we rotated the area of interest towards the region of the retina where we recorded our *t* cells (corresponding to the point (0.63, -0.35) for the left eye and (-0.63, -0.35) for the right eye view of Holmgren et al. (25)) before projecting it into equidistant coordinates. The respective videos were cropped to a total size of 1975 × 1975 *μ*m^2^ and downsampled to a pixel-size of 5 *μ*m. Furthermore, we used a copy of this dataset, but with the cricket removed from the videos. We used a total of 452 videos with a total duration of 1,464.3 seconds equal to 69,008 frames. The data was split into training (78.5%), development (15.2%), and test (6.3%) sets. In the dataset, videos were recorded in pairs for both eyes; when splitting the data, we ensured that all these pairs from individual runs were in the same split. Similarly, frames with the cricket removed were also always in the same split as their counterparts with the cricket.

#### Encoder model

We implemented the encoder as a convolutional neural network (CNN) in `tensorflow`. The CNN consisted of the following layers, resembling the vertical pathway in the retina:

1. **Input** (output shape: 395 × 395).
2. **BC spatial RFs** (output shape: 2× [111 × 111]), implemented as a 2d convolution (kernel-size: 65 × 65; stride: 3, equal to a distance of 15 *μ*m; 26).
3. **BC non-linearities** (output shape as above), implemented as a generalised sigmoid function.
4. **BC noise** (output shape as above), implemented as additive Gaussian noise with zero mean and a s.d. *σ*.
5. **RGC dendrites** (output shape: *k*× *k*; with *k* = 9 for the nasal population and *k* = 19 for the temporal population), implemented as a 3D convolution (Nasal population, kernel-size: 21 × 21; stride: 10, equal to 150 *μ*m; temporal population, kernel-size: 27 × 27; stride: 5, equal to 75 *μ*m; 22).

All encoder models we implemented shared the model parameters of the BC layers. Only the last layer, the RGC dendrites layer, differed between different encoder models.

The kernels of the BC spatial RFs layer were derived from the glutamate clusters G1 and G3 (Fig. 4) as follows. A Difference of Gaussians (DoG) was fitted to each spatial RF using the python package `Astropy` (69). During fitting, the mean and covariance matrices of the centre and surround Gaussian fits were tied, except for a linear scaling of the covariance matrix. DoG fit quality was computed as:

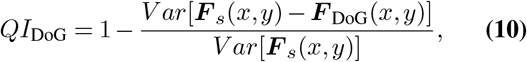

where ***F*** _DoG_(*x, y*) is the DoG fit. Finally, parameters of good fits, i.e. *QI*_DoG_ ≥ 0.35, were averaged over all ROIs per cluster and used as the BC kernels with the maximum amplitude being set to one. As non-linearities, we used the following sigmoid function:

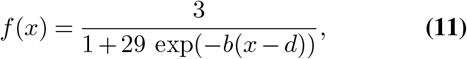

where *x* is the input and *b* and *d* are parameters per population. We defined these parameters such that the output for both groups *i* ∈ *w, s* of simulated BCs had the same response for the respective mean input 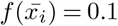per frame) and mean plus one s.d. input 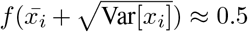 per frame). The s.d. *σ* of the additive Gaussian noise was set to 0.1. The RGC dendrites layer was implemented as two identical 2D truncated Gaussian-like functions with different weights, representing the different shares of BCs with weak and strong surround inhibition.

The parameters for the scale and cutoff were estimated as follows: First, we created a hexagonal grid representing the BC axonal terminals with a centre-to-centre spacing of 16 *μ*m (26). Then, for each morphology of *n* and *t* cells, we estimated the dendritic length per hexagon, with the central hexagon of the grid being placed on the soma (Fig. S5A). To each of these distributions, we fitted a 1D truncated Gaussian-like function (Fig. S5B), resulting in parameter estimates for the scale (Fig. S5C) and the cutoff per cell (Fig. S5D). The respective parameter means for *n* and *t* cells (Fig. S5C, D) were then used to construct the 2D RGC dendrites in the model (Fig. 5C).

#### Decoder model, training, and evaluation

For each encoder model, we created a decoder model. The decoders were implemented as ensembles of 10 CNNs with identical architectures. Each CNN consisted of 5 layers of 2D convolutions (3 filters, ReLU activation, zero padding, L_2_ regularisation with *ω* = 0.001) followed by 2D max-pooling (pool-size: 2×2, zero padding). After these layers, a dense layer (8 units, ReLU activation, L_2_ regularisation with weight *ω* = 0.003) and a single unit output layer (sigmoid activation) followed.

CNNs were randomly initialised and then trained using Adam (batch-size 16,384) to minimise the binary crossentropy loss. We used early stopping based on the validation loss that was tracked starting after 200 epochs: if the validation loss did not improve for at least 0.001 over 10 epochs, training was stopped, and the model resulting in the lowest validation loss parameters was restored. To analyse the models’ accuracies as a function of distance to cricket, we used GAMs as described above (see equation. 9), except that we used Logistic GAMs and no random effects.

## ACKNOWLEDGEMENTS

We thank Ziwei Huang for providing and explaining the code for the analysis he wrote for a previous publication. We thank Dominic Gonschorek for his feedback on the figures and manuscript. We thank Jan Lause for statistical consulting. We thank Katrin Franke, Gordon Eske and Merle Harrer for excellent technical assistance.

This work was funded by the German Research Foundation (DFG; BE 5601/6-1, BE 5601/9-1, EU 42/10-1, & EU 42/12-1), the National Natural Science Foundation of China (YR 32200810), and the Max Planck Society.

## AUTHOR CONTRIBUTIONS

Conceptualisation: JO, YR, PB, TE; Data curation: JO, YR, PS; Formal analysis: JO; Investigation: YR, JO; Methodology: JO, YR, PS; Funding acquisition: TE, PB; Project administration: TS, TE, PB; Resources: TE, PB,JK; Software: JO; Supervision: JO, TE, PB; Validation: JO; Visualisation: JO; Writing – original draft: JO, TE; Writing – review and editing: all authors.

## Supplementary Note 1: Morphology

**Fig. S1.**
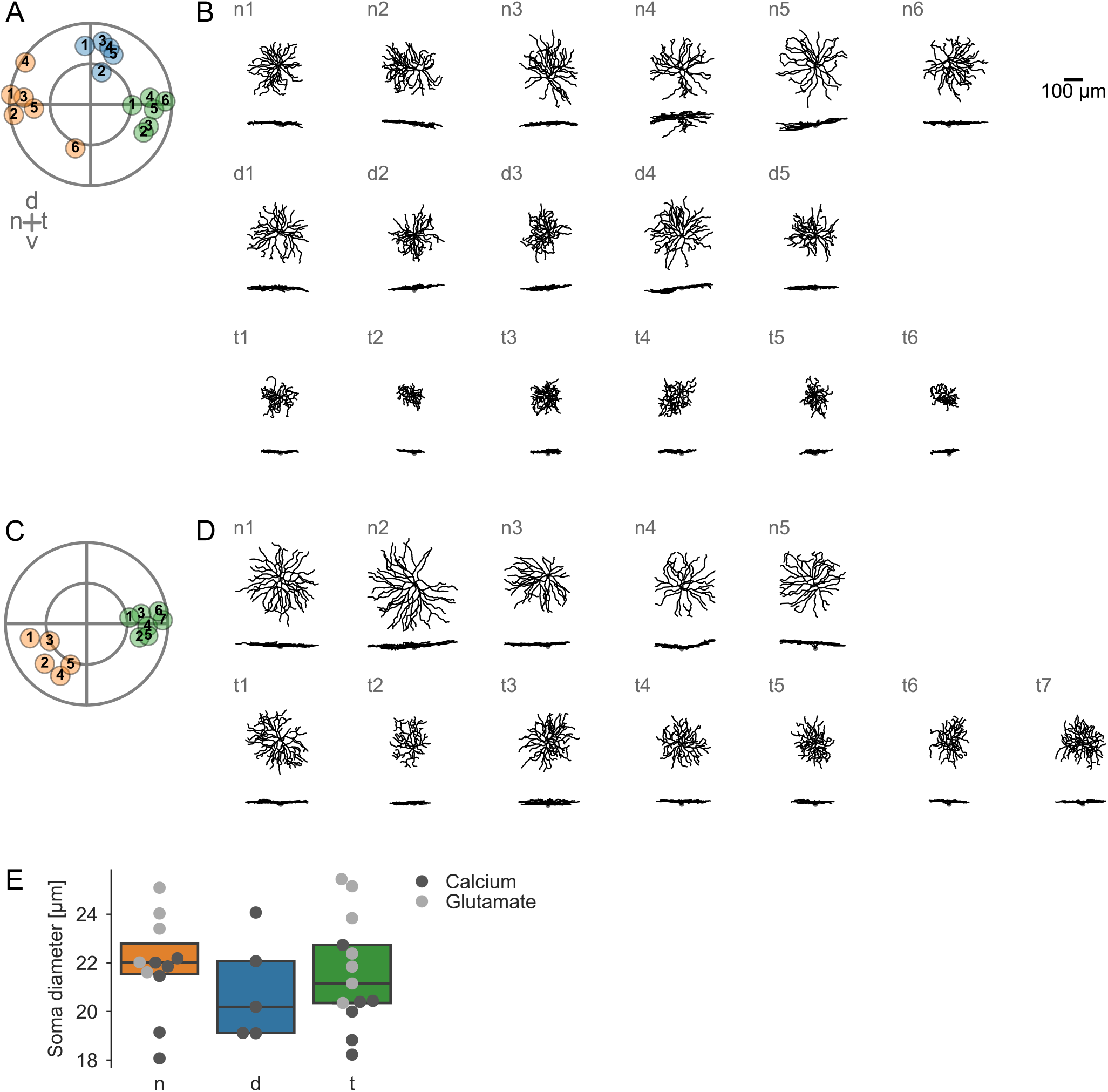
A. Cell tags of all cells with Ca^2+^recordings (*n*, nasal, orange; *d*, dorsal, blue; *t*, temporal, green). **B**. Morphologies of cell in (A). Cells are grouped by location: nasal (*n*; *top row*), dorsal (*d*; *middle row*), and temporal (*t*; *bottom row*). Within groups, cells are ordered from most nasal to most temporal. **C, D**. As in (A, B) for for the glutamate recordings. **E**. Soma diameter of cells sorted by group. Marker colour indicates if a cell is from Ca^2+^(dark grey) or glutamate (light grey) dataset.

## Supplementary Note 2: Proximal dendrite RFs

**Fig. S2.**
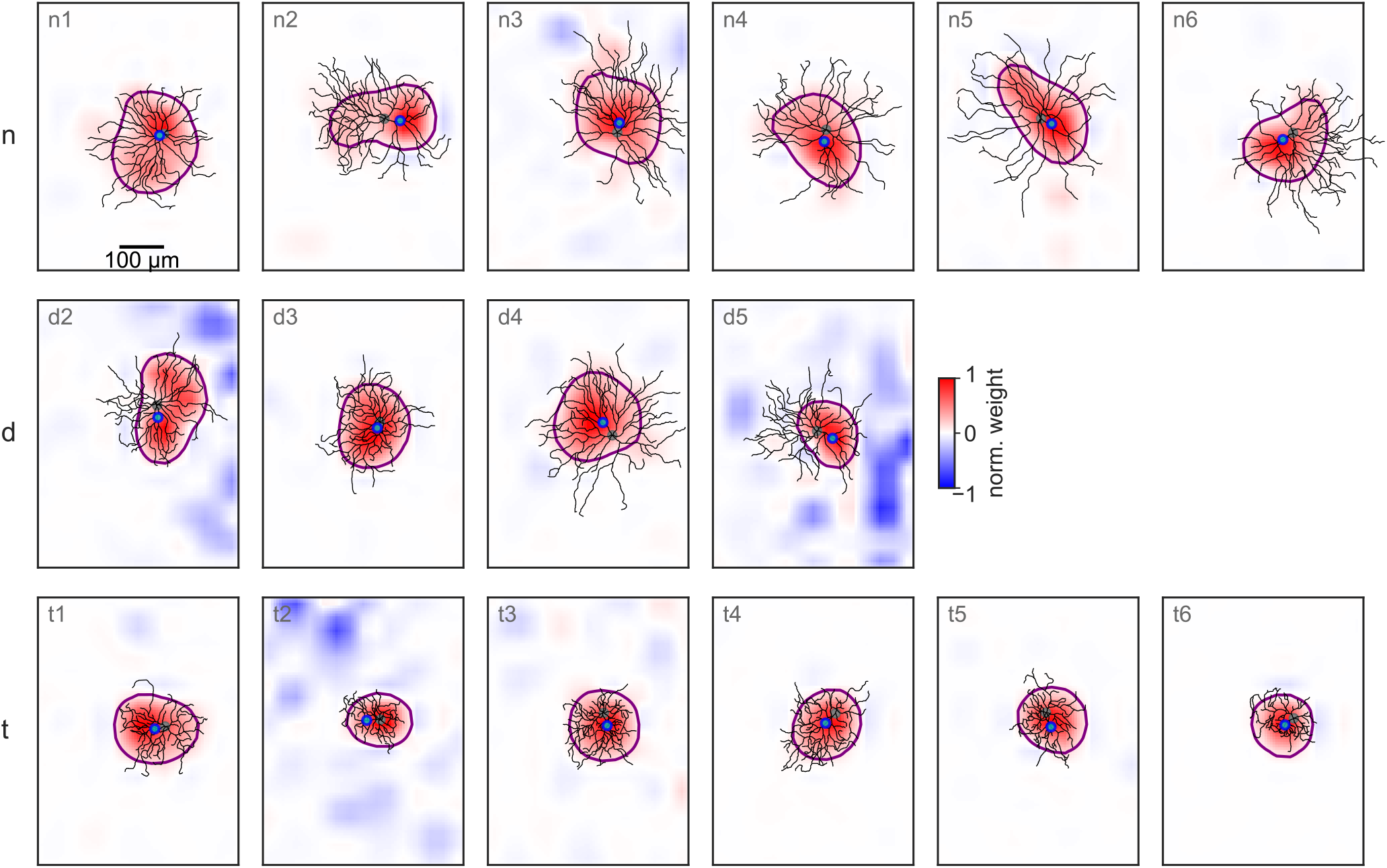
Proximal dendrite Ca^2+^RFs grouped by cell location: nasal (*n*; *top row*), dorsal (*d*; *middle row*), and temporal (*t*; *bottom row*). or exact locations, see Fig. S1A.

## Supplementary Note 3: Clustering traces

**Fig. S3.**
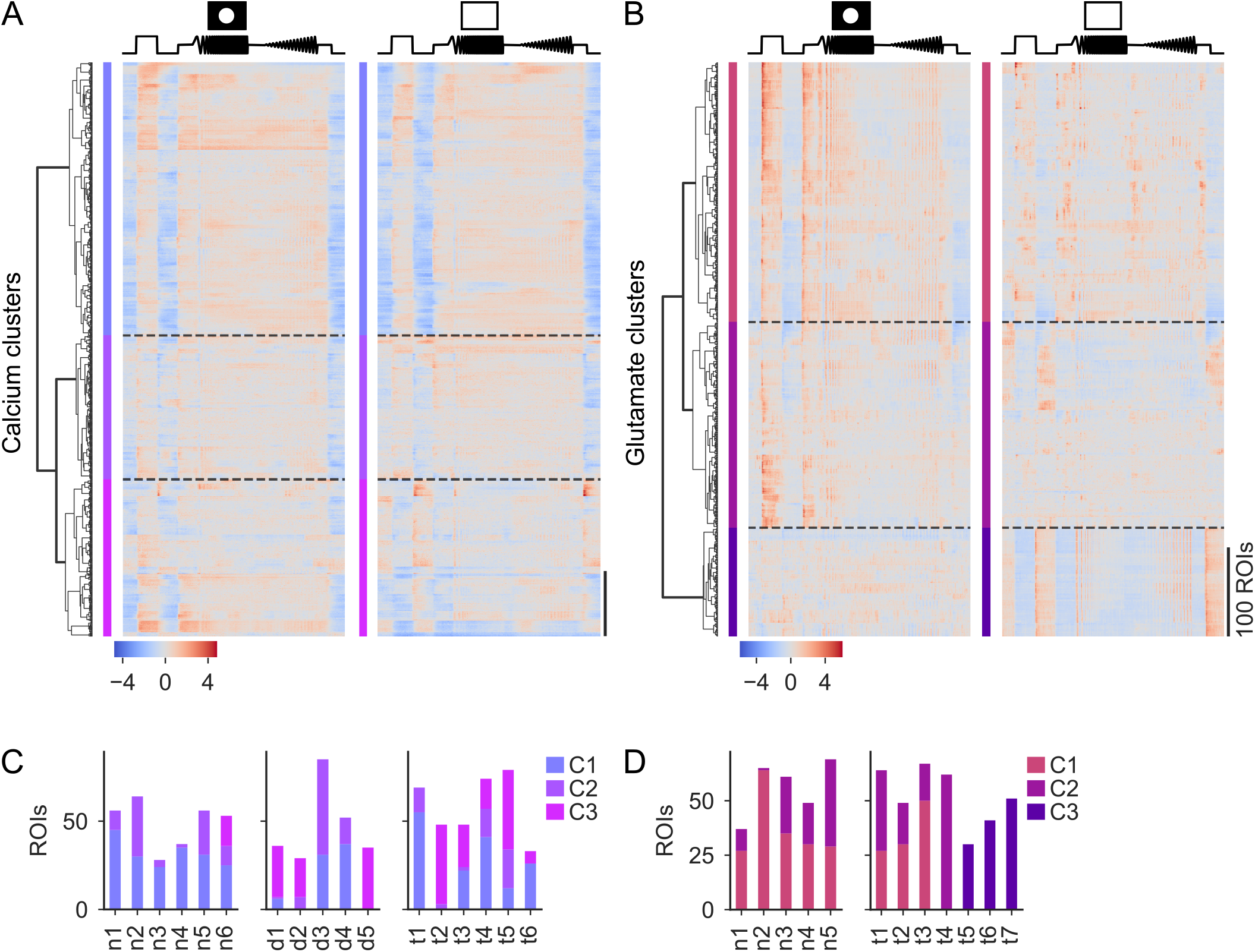
A. All chirp responses used for clustering of Ca^2+^data and the respective dendrogram. Traces are shown as heat-maps for local (left) and global (right) chirp. **B**. As in (A), but for glutamate data. **C**. Cluster counts per cell of Ca^2+^dataset. Cells are grouped by location: nasal (*n*; *left*), dorsal (*d*; *middle*), and temporal (*t*; *right*). For exact locations, see Fig. S1A. **D**. As in (C), but for glutamate data. For exact locations, see Fig. S1C.

## Supplementary Note 4: SMI-32

**Fig. S4.**
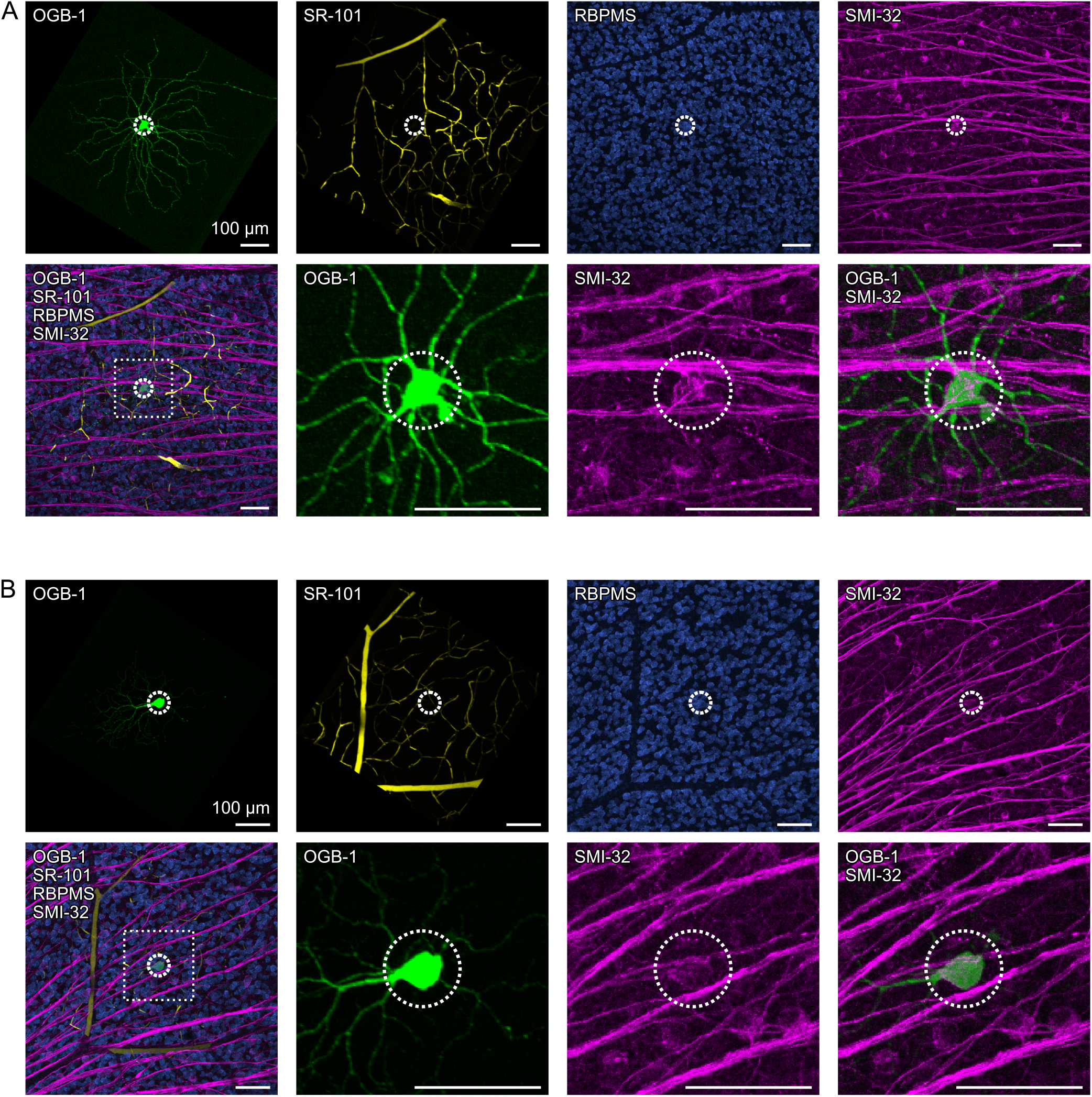
A. SMI-32 and RBPMS stainings for nasal sONα RGC. **B**. As in (A), for a temporal sONα RGC.

## Supplementary Note 5: Model parameters

**Fig. S5.**
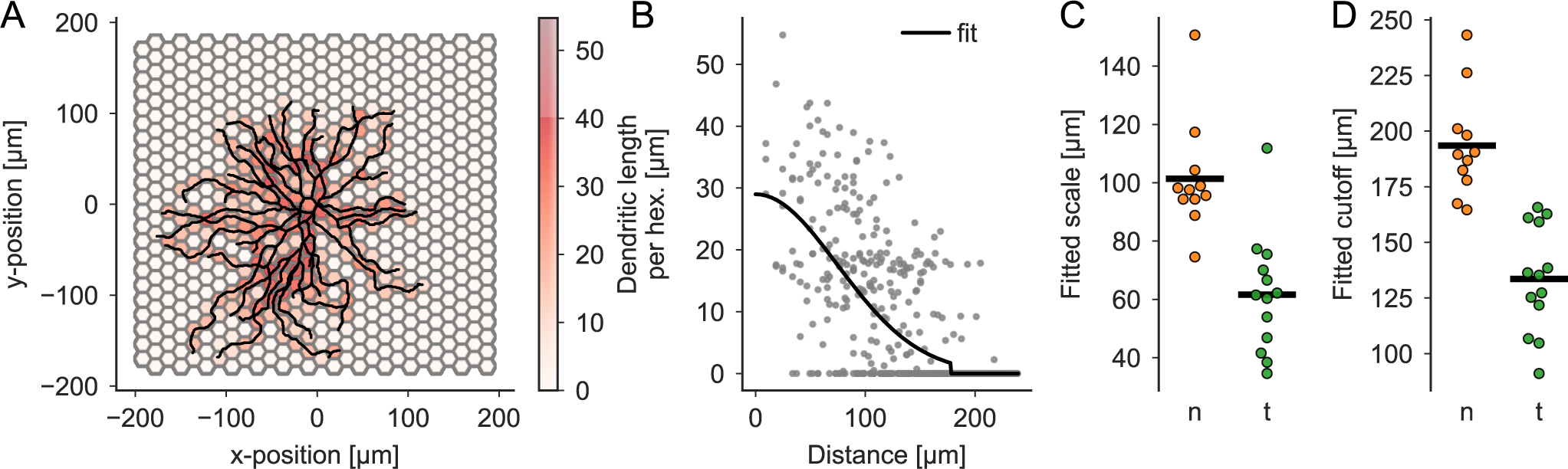
A. Dendritic densities estimated for example cell. The soma is centred on (0, 0). In each hexagon, the dendritic length is estimated (colour-coded). **B** Dendritic length as a function of distance to soma and parametric fit for cell in (A). The fitted function was a truncated bell curve, centred on zero, with an amplitude, scale and cutoff parameter optimised to fit the data. **C** Fitted scale parameter for all nasal (*n*; orange) and temporal (*t*; green) cells. **D** As in (C), but for the fitted cutoff parameter.

